# Tissue specific vulnerability to mitotic defects caused by mutations in the *Drosophila* ASPM homologue, Asp

**DOI:** 10.1101/595611

**Authors:** Lori Borgal, Margaux Quiniou, James Wakefield

## Abstract

Misregulation of candidate stem cell marker ASPM, and its *Drosophila* homologue Asp, leads to either tumour formation or microcephaly, but the functional roles contributing to each are not understood. We reverse-engineered flies to express a version of Asp (Asp^LIE^), predicted to have lost its ability to bind the phosphatase PP2A-B’. Although Asp^LIE^ flies were viable, they exhibited splayed neural stem cell spindle poles under stress, and development was substantially delayed. A tissue-level analysis of microcephaly and midgut abnormalities in Asp mutants with a compromised spindle assembly checkpoint (SAC) demonstrates tissue-specific vulnerability to mitotic defects.

## Introduction

The mitotic protein **<u>a</u>**bnormal **<u>sp</u>**indle-like **<u>m</u>**icrocephaly associated (ASPM) is a candidate stem cell marker associated with self-renewal in multiple tissues (Bikeye et al. 2010; Pai et al. 2018; Vange et al. 2015). Mutations disrupt stem cell proliferation during development (Fish et al. 2006; Marinaro et al. 2011), leading most strikingly to microcephaly (Létard et al. 2018), while increased expression enhances cancer stem cell survival (Visnyei et al. 2011; Wang et al. 2013; Pai et al. 2018; Vange et al. 2015). ASPM is a microtubule minus-end associated protein enriched at metaphase spindle poles and the midbody at telophase (Higgins et al. 2010). It contributes to mitotic processes including spindle orientation and flux, metaphase pole focusing, centriole duplication and cytokinesis (Higgins et al. 2010; Tungadi et al. 2017; Jiang et al. 2017; Jayaraman et al. 2016), although it is not yet clear which are the main contributors to stem cell dysfunction leading to microcephaly *vs.* tumour phenotypes, or how ASPM function is regulated.

Evidence supports that the *Drosophila* protein **<u>a</u>**bnormal **<u>sp</u>**indle (Asp) contributes to mitotic progression in a manner similar to its human orthologue ASPM, as it localises to spindle poles and midbody and contributes to metaphase pole focusing, spindle flux, and cytokinesis (Saunders et al. 1997; Schoborg et al. 2015; Ito and Goshima 2015; Wakefield et al. 2001). The developing larval brain of the *Drosophila* model system offers an advantage over human cell culture in the accessibility of neural stem cells to assess mitosis in a three-dimensional, tissue-specific situation. Previously, it has been shown that *asp* mutants exhibit microcephaly associated with apoptosis and neural disorganisation during development (Rujano et al. 2013). Protein phosphatase 2A (PP2A) is highly expressed in the brain, contributes to apoptosis signalling and tumourigenesis, and is a major contributor to cell cycle progression and spindle dynamics (Moura and Conde 2019; Nematullah et al. 2018). Here, we identify a potential PP2A binding site on Asp and generate animals expressing only the mutated variant. We show that, upon stress, spindles in neural progenitor cells exhibit splayed spindle poles and substantial developmental delay, but not microcephaly. Interrupting the spindle assembly checkpoint (SAC) in these flies led to pupal death associated with functional disruption in the gut rather than brain tissue, while interrupting SAC in a wild-type background caused a pupal microcephaly phenotype distinct from that in *asp* knockout flies. These data demonstrate tissue-specific differences in mitotic regulation to depend on spindle checkpoint integrity, which may constitute a basis for the stem-cell specific vulnerabilities in mitotic regulation leading to tumourigenesis or developmental disorders such as microcephaly.

## Results and Discussion

Sequence analysis of *Drosophila* Asp identified the recently described PP2A-B’ binding motif LxxIxE (Hertz et al. 2016) at the N-terminus (amino acids 350-356). This motif is also found in human ASPM, located at the C-terminus (amino acids 3266-3271); the region frequently truncated in disease (Létard et al. 2018). Of interest, the *Drosophila* sequence is directly followed by two glutamic acid residues, a characteristic described by Hertz et al (2016) to constitutively increase PP2A binding affinity. This contrasts with the human sequence which lacks these acidic residues but contains a serine in the second position that would increase binding affinity when phosphorylated, suggesting that a relationship between ASPM and human PP2A-B’ may be more heavily regulated. We generated flies expressing a version of GFP-tagged Asp (Asp^LIE^) where each key residue of the LxxIxE motif was mutated to alanine (resultant sequence AACAHA) and used these to replace endongenous Asp through rescuing *asp* null flies generated by CRISPR knockout (*asp^t25^*, Schoborg et al., 2015). GFP-expression in *asp^LIE^;asp^t25^* syncytial embryos was very similar to wild-type GFP-tagged Asp (Asp^WT^), with poleward streaming visible along metaphase spindle microtubules and a concentration of protein at the poles (Supp. Fig. 1&2).

Because the *asp* null phenotype predominantly affects neural stem cells and results in microcephaly (Schoborg et al. 2015), we investigated spindle morphology in dividing neural stem cells of *asp^LIE^;asp^t25^* larvae. Gross morphology and GFP expression of developing larval brains were similar for Asp^WT^ and Asp^LIE^ (Figure 1A). Spindle morphology was also similar at the time of dissection, but we noted that Asp^LIE^ spindles were altered as the time between dissection and fixation was increased. Systematic analyses revealed that brains fixed within 20 minutes of dissection showed no differences, while *asp^LIE^;asp^t25^* brains under dissection stress for 40 minutes prior to fixation showed splayed spindle poles and a concomitant re-distribution of GFP from a highly localised concentration at the metaphase pole to the entire spindle, when compared to *asp^WT^;asp^t25^* brains (Figures 1B-D). These data indicate that neural stem cell spindles were able to compensate *in vivo* for the *asp^LIE^* mutation but compensation was lost when brains were under stress. Recent work demonstrated that human cells expressing ASPM patient mutations had splayed spindle poles if the centrosomal protein CDKRAP5 was co-depleted and suggested that this could be a contributing factor to the microcephaly phenotype (Tungadi et al. 2017). However, although we noted that *asp^LIE^;asp^t25^* larval brains had fewer neuroblasts than their Asp^WT^-expressing counterparts at the time of dissection (Fig 2A), head size in pupae expressing Asp^LIE^ was rescued compared to the smaller head size observed in *asp* knockout pupae (Fig 4). Instead, larval development was substantially delayed in the *asp^LIE^;asp^t25^* genotype at 25°C, indicative of less efficient tissue development (Fig 2B).

**Figure 1.**
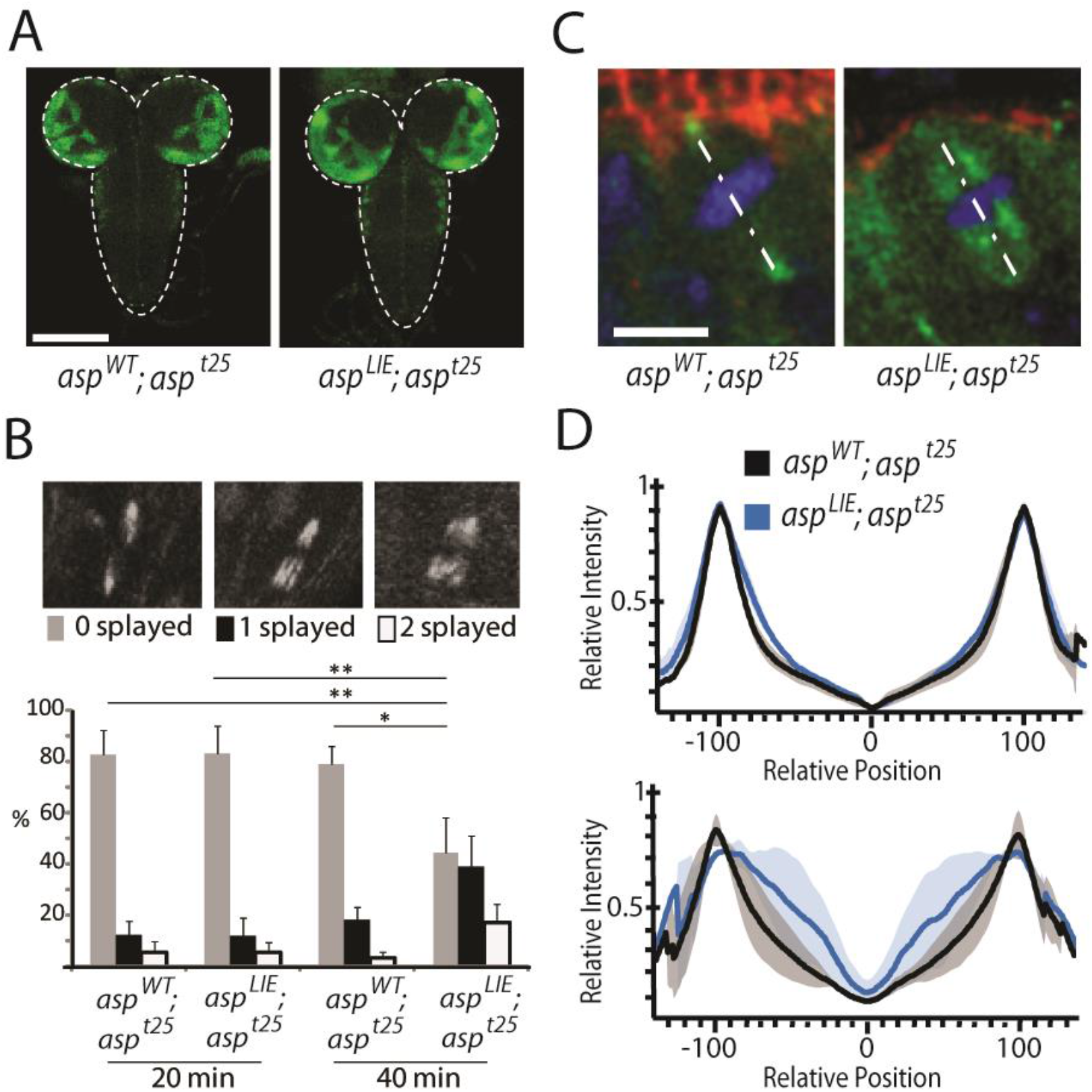
Abolishing a PP2A binding site on Asp increases neuroblast susceptibility to dissection stress. **A)** 3^rd^ instar larval brains (dashed outline). No major differences were seen in gross morphology judged by GFP expression. Scale = 150 μm. **B)** *Top.* Representative images illustrating poles scored as splayed or focused judged by GFP expression. *Bottom.* Quantification of >150 spindles scored per genotype over 3 biological replicates. Significantly fewer neuroblasts in *asp^LIE^;asp^t25^* compared to *asp^WT^;asp^t25^* larval brains had 2 focused poles after 40 minutes of dissection stress (F(3,8)=11.73, p=0.006; *=p<0.05, **=p<0.01). **C)** Example images of GFP distribution in *asp^WT^;asp^t25^ vs. asp^LIE^;asp^t25^* neural stem cell spindles at 40 minutes post dissection. Dashed line illustrates where line scan measurements were taken in D. Red = bazooka; green = Asp-GFP; blue = DAPI. Scale = 2.5 μm; maximum projection 3 μm. **D)** Quantification of GFP intensity along spindle length as indicated by dashed line in C. Peaks correspond to spindle poles. Solid line = mean; shaded area = standard deviation.

**Figure 2.**
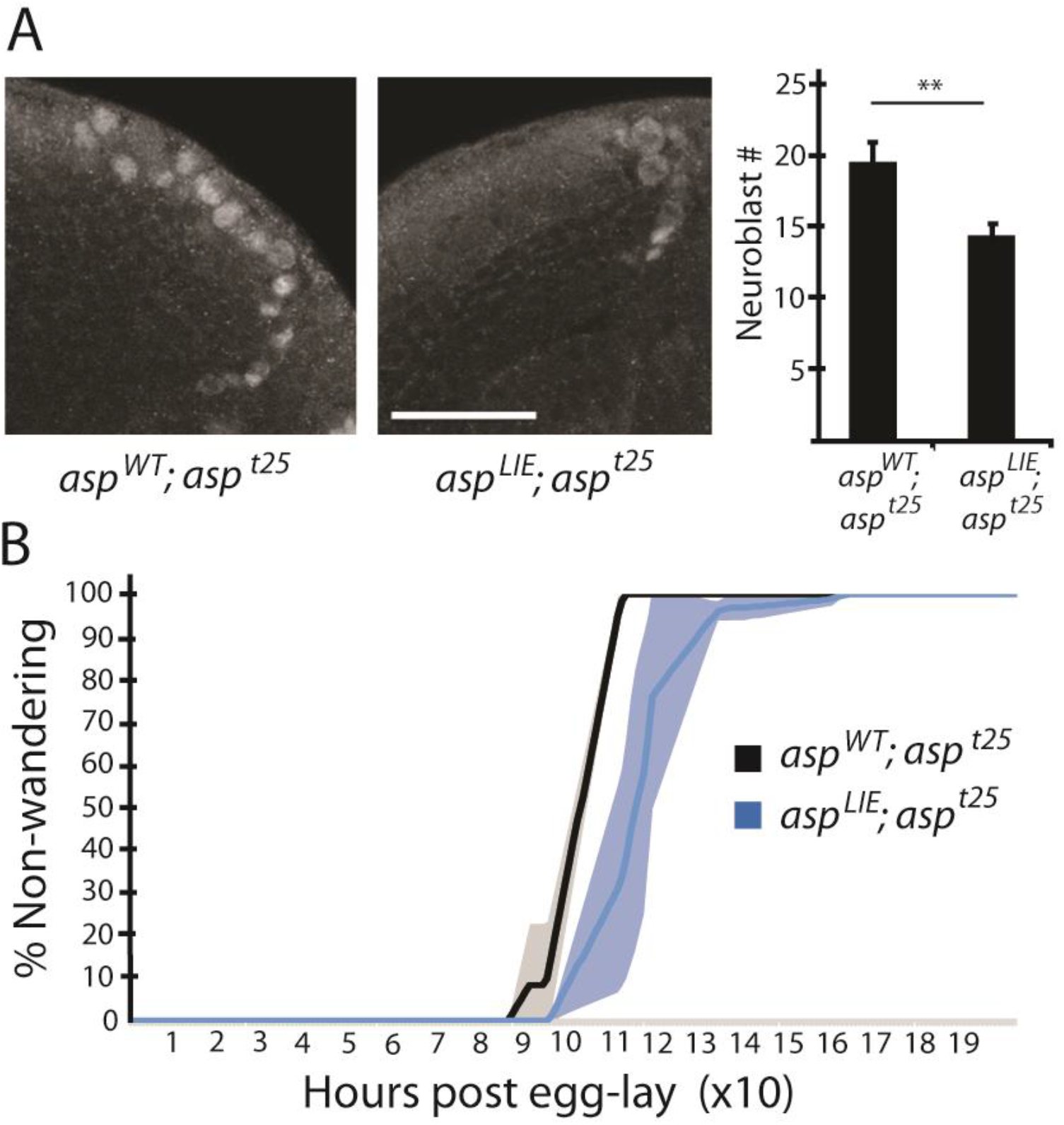
Loss of PP2A-B’ binding site on Asp yields fewer neuroblasts and developmental delay. **A)** *Left.* Cross-section of 3^rd^ instar optic lobes showing extreme examples of neuroblast depletion, immunolabelled by Deadpan in each genotype. Scale = 30 μm. *Right.* Quantification of neuroblasts per Deadpan-labelled optic lobe. Neuroblasts were scored from a single Z plane at the midpoint of each of 6 imaged lobes per genotype (t(10)=3.18, ** = p<0.01). **B)** Development of *asp^LIE^;asp^t25^* larvae was over 10 hours slower than *asp^WT^;asp^t25^*. Non-wandering larvae were scored from spiracle formation at 25°C. Solid line = mean; shaded area = standard deviation.

We wondered whether the delay in *asp^LIE^;asp^t25^* larval development reflects cellular compensation as a result of generating less efficient spindles. Cells undergoing mitosis are able to activate the spindle assembly checkpoint (SAC) and thereby delay mitotic progression until the metaphase spindle has been satisfactorily formed and all kinetochores have appropriate microtubule attachments. The gene product Mad1 is a conserved and essential SAC component previously associated with microcephaly (Poulton et al. 2017). We crossed flies harbouring a *mad1* deficiency allele (*mad1Df*) with *asp* null flies and with the *asp^LIE^;asp^t25^* engineered line, in order to reduce the capacity for SAC-mediated compensation of spindle defects in *asp^LIE^;asp^t25^* larvae. However, a developmental delay was still present in *asp^LIE^;mad1Df;asp^t25^* larvae compared to the *mad1Df* background (Fig 3A), and no microcephaly was observed (Fig 4). Strikingly though, the *asp^LIE^;mad1Df;asp^t25^* line showed significantly increased pupal death compared with either the *mad1Df* or the *asp^LIE^;asp^t25^* lines (Fig 3B). Whole body analyses revealed grossly distended pupal midguts (Fig 3C), phenocopying the visceral myopathy associated peristalsis deficit observed upon reduced expression of the spindle-associated *misato* (Min et al. 2017). In humans and flies, the gut visceral muscle serves as a niche for intestinal stem cells (Min et al. 2017; Biteau and Jasper 2011). These data demonstrate differences in tissue specific robustness to multiple mitosis-related challenges: loss of the SAC in gut tissue sensitizes it to Asp mis-regulation through loss of the PP2A binding site, while brain tissue is still able to compensate.

**Figure 3.**
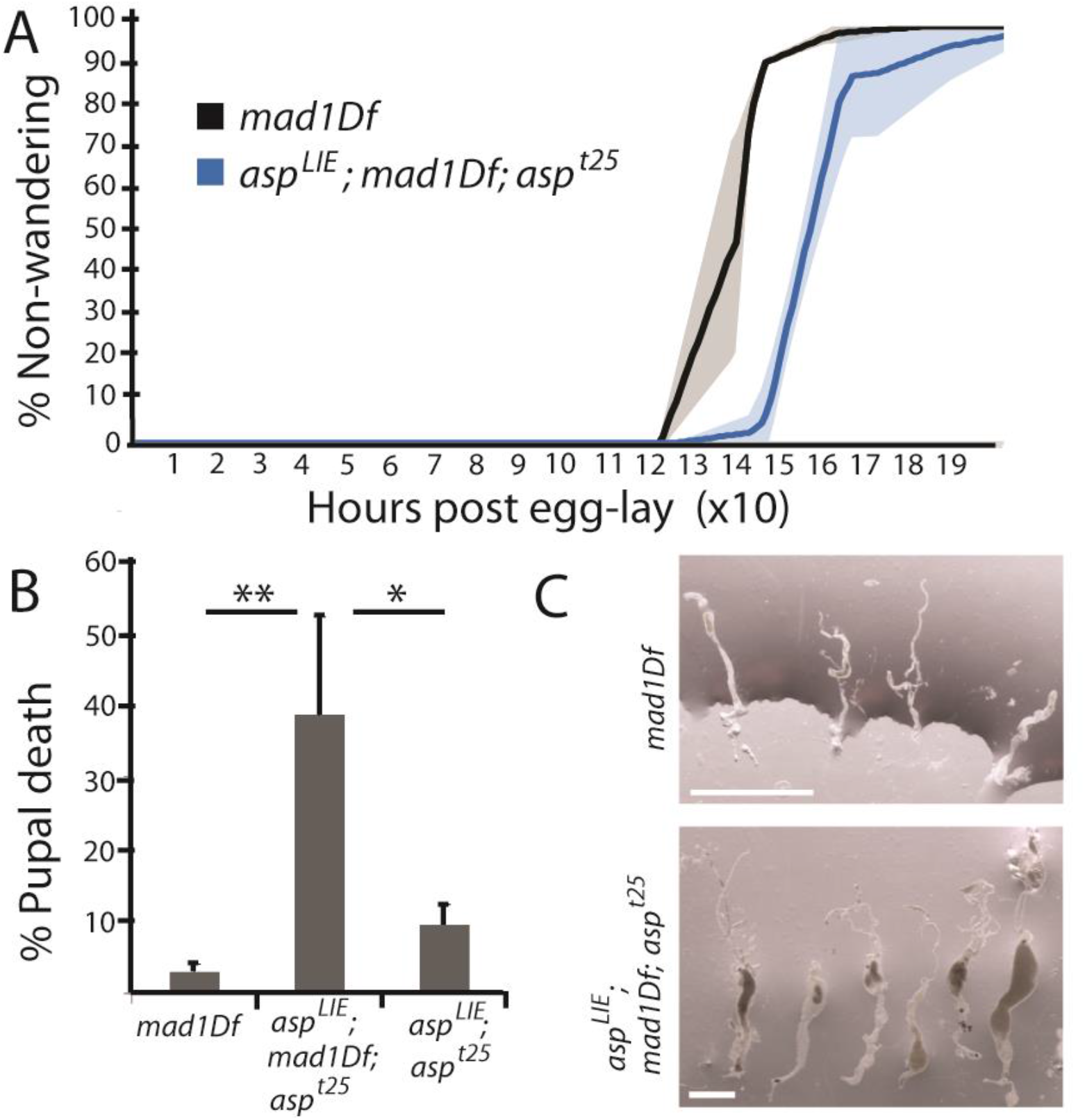
Reducing SAC efficiency does not abrogate developmental delay caused by PP2A-B’ binding site mutation, but does increase pupal death. **A)** Development of *asp^LIE^; mad1Df; asp^t25^* larvae was over 10 hours slower than in the *mad1Df* background. Non-motile larvae were scored from spiracle formation at 18°C. Solid line = mean; shaded area = standard deviation. **B)** Failure to eclose was significantly increased in *asp^LIE^; mad1Df; asp^t25^* compared to either of the *asp^LIE^; asp^t25^* or *mad1Df* backgrounds. Failure was scored if pupae with dark wings had not eclosed after 24 hours (F(2,10)=10.12, p=0.004; *=p<0.05, ** =p<0.01). **C)** Dissected midguts of *asp^LIE^; mad1Df; asp^t25^* pupae that had failed to eclose were often distended. This was never observed in the *mad1Df* background with unmodified Asp. Scale = 500 μm.

**Figure 4.**
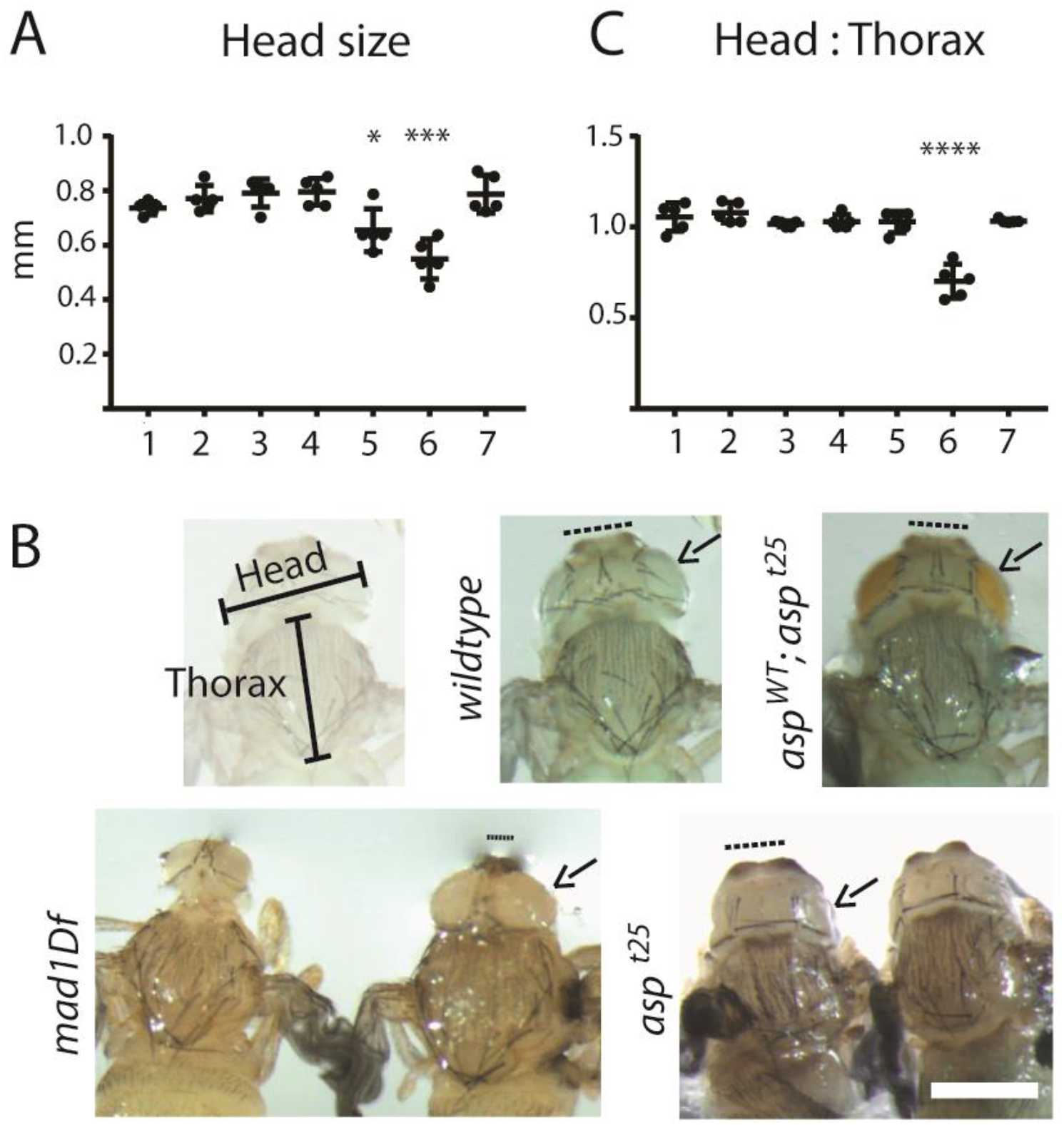
Microcephaly manifests differently after disruption of *asp vs.* SAC. **A)** Head size in millimetres for genotypes as follows: 1) *wildtype* 2) *asp^WT^;asp^t25^* 3) *asp^LIE^;asp^t25^* 4) *asp^LIE^;mad1Df; asp^t25^* 5) *asp^t25^* 6) *mad1Df* 7) *mad1Df; asp^t25^.* Head sizes were measured for slow developers (13 days post egg lay) that failed to eclose 24 hours after wings were black. Mean head size for *mad1Df* pupae was significantly smaller than all other genotypes. Mean head size for *asp^t25^* pupae was significantly smaller than *asp^LIE^;asp^t25^* pupae, *asp^LIE^;mad1Df;asp^t25^* pupae, and *mad1Df; asp^2^* pupae (F(6, 28) = 12.38, p<0.0001; * =p<0.05; *** =p<0.005). **B)** Images of head and thorax for the indicated genotypes, and schematic illustrating measurement strategy. Note eye sizes (arrows) compared to central brain (dashed line). Scale = 500 μm. **C)** Ratio of head: thorax measurements for genotypes labelled as in A. This ratio consistently approached 1:1 except for *mad1Df* pupae where head size was significantly less than thorax (F(6, 28) = 25.36, p<0.0001; **** = p<0.0001).

In addition to differences in *asp^LIE^* brain *vs*. gut tissue susceptibility to *mad1* interruption, we observed *mad1*-dependent differences within central brain *versus* optic lobe microcephaly. Small head size is well documented to be a consequence of *asp* loss (Rujano et al. 2013; Schoborg et al. 2015) and has also been reported to result from SAC disruption caused by Mad1 deficiency (Poulton et al. 2017). Our data supports this, with pharate head size significantly reduced in both *asp^t25^* and *mad1Df* lines (Fig 4A). However, visual comparison revealed a stark difference in the manifestation of the microcephaly phenotype. In agreement with Rujano et al. (2013), pharate head size resulting from lack of Asp expression was primarily reduced in optic lobes, with very little decrease in central brain size (Fig 4B). In contrast, there was a dramatic head size reduction in the central brain of *mad1Df* pupae, with no apparent decrease in eye size. It is important to note that for both genotypes there was a high consistency of microcephaly observed upon dissection of slow-developing pupae that had failed to eclose, but a much greater variability ranging to normal in adults, indicating that compensation during development exists in both cases. Furthermore, although the *asp^t25^* knockout brains had disproportionately reduced optic lobe size, the overall head size reduction was in proportion to an overall body size reduction (Fig 4C). This indicates that in addition to a specific susceptibility to reduced functional Asp in developing optic lobes, there is a general susceptibility across all tissues. Similarly, while the most recent clinical analysis shows a more profound reduction in head size of ASPM-microcephaly patients at birth, body weight was also documented as an average of 1.2 standard deviations smaller than the mean population control (Létard et al. 2018), indicating that there may also be a milder developmental phenotype across multiple tissues in humans. Finally, no microcephaly was observed in the *mad1Df; asp^t25^* genotype. This supports, on the one hand, that reduced SAC efficiency ameliorates *asp*-induced microcephaly, and demonstrates on the other hand that the central brain microcephaly observed due to *mad1* deficit is dependent, at least in part, on Asp function. Indeed, this genotype was healthy at 25°C, whereas *mad1* deficient flies in a wildtype Asp background required development to be slowed at 18°C for survival. Furthermore, homozygote *mad1Df* flies were observed in an *asp* knockout background when housed at 18°C; these were never recovered in a wildtype *asp* background. This data is reflective of *mad2* spindle assembly checkpoint mutants restoring mitotic index to wildtype levels when crossed with *asp* mutants (Buffin et al. 2007).

These data demonstrate tissue specific susceptibility to disrupted metaphase spindle formation. In mammals, the loss of ASPM function has been thought to specifically impede neurogenesis because the specialized cellular morphology of neural progenitors renders them extremely susceptible to alterations in spindle orientation (Fish et al. 2006; Gai et al. 2016). However, two recent studies have provided evidence uncoupling spindle orientation from the microcephaly phenotype, pointing instead to apoptosis (Insolera et al. 2014) or impaired centriole duplication (Jayaraman et al. 2016). In *Drosophila*, apoptosis has been associated with severe disorganisation in third instar larval brains and microcephaly (Rujano et al. 2013). This developmental stage coincides with increased mitosis after a period of dormancy (Poulton et al. 2017). Mitotic errors can lead to aneuploidy, which is well tolerated in mammalian and insect neural progenitors (Damiani et al. 2016; Poulton et al. 2017) but may be associated with tumourigenesis in other cell types (Simonetti et al. 2019). Cells that cannot force mitotic completion in this way can either exit the cell cycle as a 4N cell or enter apoptosis. The winner between these outcomes may be mediated in part by the relative activity of the SAC.

Previous data demonstrated that *Drosophila* neural development was unaltered by loss of the SAC, but did report sterility in the *mad2-mad2B* double mutants (Buffin et al. 2007), highlighting a tissue specific vulnerability to loss of the SAC as was seen in *asp^LIE^;mad1Df;asp^t25^* midgut tissue in the present study. The authors demonstrated that the SAC becomes critical in *Drosophila* when mitosis is rendered less efficient (Buffin et al. 2007). Loss of *mad1* has been reported to impede kinetochore-microtubule attachment, which combined with SAC disruption may cause a more striking phenotype than observed in *mad2* mutants (Emre et al. 2011). Alternatively, identifying mitotic abnormalities in *mad2* mutants may require observing slow developers or parsing the dataset by neuroblast subtype, as the current data demonstrates a difference in vulnerability to *mad1* loss between central brain and optic lobe. Differences in SAC importance between cell types may reflect the speed with which mitosis is required to occur: it has been proposed that SAC activation in stem cells may be required to prolong mitosis allowing time for correct spindle alignment (Mantel and Broxmeyer 2007). Neuroblasts can establish spindle polarity despite complete centrosome detachment (Schoborg et al. 2015), indicating that in this cell type spindle assembly may rely primarily on chromatin-nucleated microtubules. This pathway is slower to assemble spindles post nuclear envelope breakdown (Hayward et al. 2014), and these cells may require a different baseline of SAC activity than their neighbours. Indeed, aneuploidy *vs.* apoptosis were demonstrated to be tissue-specific responses to loss of centrosomes, which primed a tissue-specific response to SAC disruption (Poulton et al. 2017). Any perturbation in mitotic efficiency, such as impediments to spindle pole formation or timely centriole duplication, may activate the SAC with varying consequences dependent upon cell type. The present work underscores the importance of considering baseline dynamics for spindle formation and spindle checkpoint when investigating development and disease susceptibility in different cell types and tissues.

## Materials and Methods

### Fly lines and cloning

Flies were housed according to standard procedures at room temperature except during assays as indicated in text. The *asp^WT^* and *asp^t25^* lines were gifts from Nasser Rusan (Schoborg et al. 2015). The *msd1Df* line was purchased from Bloomington (Stock 4966; Df(2R)w45-30n cn1; deletes mad1, Poulton et al. 2017). The *asp^LIE^* line was generated by Quikchange PCR of the pENTR vector containing *asp^WT^* cDNA (gift from Nasser Rusan (Schoborg et al. 2015), sequence validated and recombined to a Ubq plasmid (gift from Renata Basto (Peel et al. 2007)), then injected into w^1118^ embryos by BestGene Inc. (Chino Hills, CA, USA). The wildtype line (Stock 416; w[1118]; Df(3L)GN19/TM3, ry[*] su(Hw)[2] Sb[1]) was obtained from Bloomington.

### Larval brain dissection and immunofluorescence

Brains were dissected from third instar larvae at 96 hours post egg lay and fixed for 20 minutes in 4% Formaldehyde in PBS with 1 mM EGTA and 10 mM MgCl_2_. Brains were washed in PBS plus 1% Triton X-100, blocked in 3% BSA, incubated with primary antibody overnight followed by washing and incubation with species-appropriate secondary antibody at 1:200 (Alexa Fluor 555, Invitrogen) for at least 2 hours, washing and incubation in 50% glycerol for 2 hours, then 70% glycerol overnight. The GFP signal was retained through this process. Primary antibodies were as follows: anti-deadpan (guinea pig), used 1:3000, gift from Christian Q Doe; and anti-Bazooka (rabbit), used 1:1000, gift from Andreas Wodarz. Brains were mounted in 35% glycerol, 50% Vectashield Antifade Mounting Medium with DAPI (Vector Laboratories, UK) in PBS, mounted inside reinforcement rings and coverslipped for imaging.

### Development Assays

Larval development was assessed at 25 °C for *asp^LIE^;asp^t25^* and *asp^WT^;asp^t25^*, and at room temperature for *mad1Df* and *asp^LIE^; mad1Df; asp^t25^.* Adult flies were left to egg-lay in tubes for 1h at 25 °C or 3h at room temperature. The number of non-wandering 3^rd^ instar larvae was quantified at regular time intervals until 8 days post egg-lay (192 HPE) to measure developmental time course until pupation. Pupal death was scored when mature pupae with visible dark wings had not eclosed 24 hours later. Results were analysed by one-way ANOVA and Tukey’s post-hoc.

### Neural Stem Cell Analyses

Imaging was performed using a Leica SP8 confocal laser-scanning microscope with Leica Application Suite X. For each experiment, at least 3 larval brains were imaged per strain. Z-stacks of optic lobe neuroblasts were taken. All image processing was performed using FIJI (ImageJ Inc.(Schindelin et al. 2012)). For all metaphase spindles detected within each z-stack, the planes which comprised the spindle were selected and projected into a single maximum-intensity image. Metaphase spindle pole splaying was scored as positive or negative by a blinded experimenter according to the Asp-GFP signal. Ambiguous spindles were excluded from analysis (13 removed out of 609 total). Results of 3 separate experiments run were analysed for bi-polar focused spindles only by one-way repeated measures ANOVA with Tukey’s post-hoc. Asp distribution along spindles was measured using the Line Selection tool in FIJI (ImageJ Inc.) to draw a line intersecting both spindle poles, which was then measured for fluorescence intensity using the Plot Profile function. The lowest intensity recorded in each line scan was considered as background signal and removed. Intensities were then normalised on a scale from 0 to 1. For all intensity curves, distance normalisation was performed in order to align the 2 highest fluorescence peaks (corresponding to the spindle poles). A Matlab program was created to interpolate all intensity curves, which allowed the mean fluorescence intensity and standard deviation to be calculated along the length of the spindle. Neuroblasts were counted as Deadpan positive cells in optic lobes of 3^rd^ instar larval brains. After brain lobe imaging at low magnification (20x objective), the slice presenting the largest brain lobe diameter was selected for analysis, ensuring this single slice reflected the mid-section of the sample. Six lobes from three brains were quantified per genotype. Results were analysed by one-way ANOVA with Tukey’s post-hoc.

### Pharate head size and midgut dissection

Eggs were laid overnight in standard housing and kept at room temperature (21°C). Pharates that were slow to develop had black wings visible on day 13. Those that had not eclosed 24 hours later were removed from the pupa manually in PBS. Head and thorax were measured by calibrated graticule. Midguts were examined by manually removing from abdomen. Images were taken with Olympus SZX16 stereoscope and Cell^D software (Olympus Soft Imaging Solutions GmBH). Data was analysed by one-way ANOVA with Tukey’s post hoc.

## Acknowledgements

The authors would like to thank members of the Wakefield lab and Living Systems Institute for helpful discussion.

